# Cross-talk between m6A and m1A regulators, YTHDF2 and ALKBH3 fine-tunes mRNA expression

**DOI:** 10.1101/589747

**Authors:** Nga Lao, Niall Barron

## Abstract

The recently re-discovered interest in N^6^-methyl adenosine (m6A) - one of more than a hundred modifications found on eukaryotic mRNA (known as epi-transcriptomic codes) - is currently one of the most topical areas in science. The m6A methylation impacts on all aspects of cellular RNA metabolism and several physiological processes. Although less abundant than the m6A epitranscriptomic mark, the m1A methylation has recently also attracted interest due to its dynamic nature in response to physiological changes. We investigated the role of the YTH domain-containing m6A reader protein family and the m1A eraser ALKBH3 on the expression of a transgene in mammalian cells. We present the first evidence that expression of a transgene is subjected to co-ordinated regulation by both m6A and m1A regulators. In addition, we provide genetic data implicating that the m6A reader YTHDF2 can read m1A site. Furthermore, we show that the m1A eraser ALKBH3 is a target of m1A methylation.

## INTRODUCTION

A large number of studies in the last 6 years have convincingly proven the impact of N^6^-methyl adenosine (m6A) - the most abundant modification on eukaryotic mRNA - on all aspects of RNA metabolism including spicing, stability, translation, microRNA processing (1–7) and several physiological processes including cancer, immunity and memory (8–10). The reversible m6A modification is installed by the m6A writer complex (11) containing the catalytic core METTL3 and METTL14 proteins (12) and removed by the m6A erasers ALKBH5 and FTO (13, 14). The m6A readers bind m6A RNA targets to mediate their fate. At present, more than twenty m6A have been identified (1, 6, 15–17), of which, the 5-member YT521-B Homology (YTH) domain-containing protein reader family (YTHDF1-3, YTHDC1, and YTHDC2), which directly bind m6A through the YTH domain, are the most studied (18). Notably, the three cytoplasmic YTHDF members (YHTDF1-3 or DF1-3 for short) have been shown to act in an integrated and coordinated manner in affecting the fate of their shared m6A RNA targets and ultimately protein expression. Cooperation between DF1 and DF3 has been shown to promote translation, whereas DF2 and DF3 conspire to accelerate m6A-containing RNA decay (19, 20).

N^1^-methyl adenosine (m1A) is another epigenetic mark on mRNA that has recently attracted interest due to the observation that it is dynamically regulated in response to changes in cellular or physiological conditions, and is generally positively correlated with protein expression (21–23). The tRNA methyltransferase complex TRMT6/61A has been implicated in m1A modification of some nuclear mRNAs at m1A sites with t-RNA T-loop-like structures (23, 24), however, for the majority of m1A sites in mRNA, the responsible m1A writers are yet to be identified. ALKBH3 - also known as prostate cancer antigen-1 (PCA-1) - has been shown to be an m1A eraser in mammalian cells (21, 22). Given the abundance of these modifications in mRNA, we were interested in establishing if the expression of a transgene could be impacted by m6A and m1A regulation in mammalian cells used for commercial production of biopharmaceutical proteins – with a view to increasing the efficiency of therapeutic protein production. While high transcription rates are typically achievable from modern expression vectors, this doesn’t always translate into high protein yields. We reasoned that further improving productivity in mammalian cells might be achieved by modulating the regulators of these epitranscriptomic marks which have shown striking impact in basic cellular mRNA metabolism and translation. However, knowing with certainty if a transgene is subjected to m6A or m1A regulation remains a challenge at present, as m6A methylation also occurs in sequences outside the canonical DRACH motif (D=A/G/U, R=A/G, A=m6A, and H=A/C/U), and not all DRACH motifs are m6A methylated (25). In addition, a universal method to precisely identify m1A in mRNA is not yet available (26). In this paper, we characterised the impact of silencing of the m6A readers DF1-3 and m1A eraser ALKBH3 on the expression of reporter Green Fluorescence Protein (GFP) gene in Chinese Hamster Ovary (CHO) cells which are the most commonly used cells for industrial production of pharmaceutical proteins. We demonstrate that the cross-talk between m6A and m1A regulators plays an important role in the expression of a transgene in mammalian cells.

## MATERIAL AND METHODS

### Cell lines and reagents

CHO-K1 cell line (ATCC^®^ CCL-61) was adapted for suspension culture and routinely tested for mycoplasma contamination. Cells were maintained in Serum Free Media (SFM) supplemented with 2.52g/L anti-clumping polyvinyl alcohol (PVA) unless otherwise stated. Cultures were grown in a Climo-Shaker (Kühner) at 37°C, 80% humidity and 5% CO2, 170 rpm. GFP and both cell growth and viability were monitored using the Guava Express Plus and Guava Pro programmes, respectively on the Guava EasyCyte (Merck Millipore). Primary antibodies used in the experiments are listed as follows; anti-GAPDH (60004-1-Ig, Proteintech), anti-m1A mAb (D345-3, Medical & Biological Laboratories CO., LTD), anti-m6A (E1610S, NEB; 202003, Synaptic Systems), Flag tag Monoclonal Antibody (66008-2-Ig, Proteintech); anti-human EPO (AB-286-NA; R&D Systems). Secondary antibodies: IRDye^®^ 800CW Goat anti-Mouse IgG (P/N 925-32210), IRDye^®^ 680LT Goat anti-MouseIgG (P/N 925-68020) from LI-COR. Protein A/G Plus-Agarose (sc-2003; Santa Cruz Biotechnology), RNase Inhibitor (N8080119, Applied Biosystems™). Standard EPO (329871, Merc Millipore). TransIT-X2^®^ transfection reagent (Mĭrus) was used for transfection. DNA free total and mRNA were prepared using Direct-zol RNA kit (R2060, Zymo Research) and GenElute™ mRNA Miniprep kit (MRN10, SIGMA-ALDRICH), respectively.

### Vectors

The vector CMV-d2GFP-HYG (27) which contains a coding sequence for the unstable d2GFP between the CMV promoter and polyA was used in this work. Clone OHu31338D containing the open reading frame of the human ALKBH3 in pcDNA3.1 with C-terminal FLAG tag was obtained from GenScript, U.S.A. The overexpression vector CMV-DF2 was constructed as follows: The open reading frame (ORF) of DF2 (GenBank accession number XM_007626785) was first amplified using cDNA made from DNA-free total RNA from CHO-K1 as a template and primer pair DF2for and DF2rev. The resulting amplicon was used as a template to incorporate 3 FLAG epitopes upstream of the start codon using overlapping forward primers (FLAG2YN and 3FLAGBamfor) and reverse primer Y2endXho. The resulting fragment was cloned into pcDNA3 at the BamHI and XhoI sites. Primer sequences are listed in Table S1A.

### Transfection

Transfection was carried out in a 24 well suspension plate using TransIT-X2^®^ transfection reagent, following the protocol recommended by the manufacturer (Mĭrus). For co-transfection of siRNA and plasmid DNA, the siRNA was transfected 16-24 hours in advance of the plasmids. The final concentration of siRNA was 25nM and 400ng DNA plasmid per 1×10^6^ cells/mL.SiRNA was designed and supplied by Integrated DNA Technologies (IDT). The siNC was also bought from IDT. SiRNA sequences are listed in Table S1B.

### RT-PCR

cDNA was prepared using High Capacity cDNA Reverse Transcription Kit (4368814, Applied Biosystems) following the manufacturer’s protocol. RT-PCR was performed using FAST SYBR kit (4385616, Applied Biosystems) with three technical replicates and analyzed on an Applied Biosystems 7500 FAST and Real-Time PCR system. The fold change of gene expression was calculated using the 2^(-ΔΔCt) method. Primers used in qPCR are listed in Table S1C.

### Western blot analysis

Cells were lysed in RIPA buffer (R0278, SIGMA-ALDRICH) containing Halt™ protease inhibitor cocktail (87786, Thermo Scientific) at 4°C for 1 hour with gentle rotation. Cell debris was removed by centrifugation at 14,324g, protein lysate was heated at 70oC for 10min in the presence of 1X loading buffer. Proteins were separated on a precast 4-12%Bis-Tris Plus gel in Bolt™ MOPS SDS running buffer (Invitrogen). Gels were blotted on a Nitrocellulose membrane (Amersham TM Protran™ 0.45um, GE Healthcare Life Science) using Thermo Scientific Pierce Power Blotter (# 22834) in Pierce 1 step transfer buffer (84731, Thermo Scientific). Blots were scanned and quantitatively analyzed using the Odyssey ^®^ Fc Imaging System.

### M6A and m1A immunoprecipitation

M6A and m1A immunoprecipitation were adapted from the protocols provided in the EpiMark^®^ N6-Methyladenosine Enrichment Kit (NEB #E1610S, New England BioLabs Inc) and that previously described (28). Briefly, 3.5ug mRNA was incubated for 3 hours with Protein A/G agarose beads that were pre-incubated with 5ug of anti-m6A, -m1A or -Gapdh antibody for 1 hour with head to tail rotation in reaction buffer (150 mM NaCl, 10 mM Tris-HCl, pH 7.5, 0.1% NP-40). Beads were washed twice with reaction buffer, twice in low salt buffer (50 mM NaCl, 10 mM Tris-HCl, pH 7.5, 0.1% NP-40) and twice in high salt buffer (500 mM NaCl, 10 mM Tris-HCl, pH 7.5, 0.1% NP-40). Immunoprecipitated RNA was eluted using reaction buffer containing 6.7mM N6-methyladenosine (00587, Chem_Impex International, Inc., U.S.A.) or 3mg/mL N1-methyladenosine (sc-216121, Santa Cruz Biotechnology). All buffers contained RNase inhibitor. 350uL of eluted RNA was precipitated with 2.5 volume of absolute ethanol, 0.3M NaAc, pH5.5 and 1.5uL glycogen (G1767, SIGMA Aldrich) overnight at −80°C, washed twice in 75% Ethanol and resuspended in 15uL water. cDNA synthesis and qPCR were carried out as described in EpiMark^®^ N6-Methyladenosine Enrichment Kit using High-Capacity cDNA synthesis and Fast SYBR™ Green Master Mix (4368814 and 4385616, Applied Biosystems™) kits. The percentage of immunoprecipitate (IP) to Input was calculated as described in the Magna MeRIPTM m6A kit (17-10499, Millipore). mRNA controls (m6A RNA and unmodified RNA) were used as described in the EpiMark^®^ kit.

## RESULTS AND DISCUSSION

### GFP expression is affected by both m6A and m1A regulators

First, we found that there were no m6A sites predicted in the GFP transcript using on-line prediction software (http://www.cuilab.cn/sramp/) (29) which is based on the presence of the DRACH consensus motif. However, there are many GAC motifs which have been shown to be a preferable target for the m6A reader DF1/2 (7, 30), two of which are GACT. In addition, we found two predicted sites for m1A within the coding sequence of GFP, one of which (GTTCGA) was previously identified by two studies mapping m1A occurrence in human cells (23, 24) and a second putative m1A site (GGTGGA) identical to that present on the human SRSF gene (24) (Figure 1A). We characterized the expression of GFP in CHO-K1 cells co-transfected with siRNA targeting various m6A writers; the catalytic core METTL3, METTL14, and a regulator of writer complex formation (WTAP), m6A erasers (ALKBH5 and FTO) and the m1A eraser ALKBH3. We found that the level of GFP protein expression was substantially reduced when the m6A readers DF1 or DF3 were silenced, as opposed to silencing DF2 which increased GFP expression nearly two-fold (Figure 1B). The relative levels of GFP mRNA were 4-fold higher in DF2-silenced cells compared to that of the siRNA negative control (NC) suggesting that the GFP mRNA was more stable when DF2 was depleted (Figure 1B), consistent with DF2’s putative role in accelerating mRNA degradation (7). This is not unexpected, as each of the YTH domain containing proteins DF1-3 from CHO shares at least 94% amino acids identity with their mammalian orthologs including essential residues for binding to m6A (Figure S1). A pronounced impact of DF2 knockdown was also observed on the expression of human Erythropoietin (EPO), a drug used for the treatment of some forms of anemia (Figure S2). The expression of GFP was reduced by nearly 40% in siALKBH3 transfected cells while only being negligibly affected when other m6A regulators (METTL3, METTL14, and WTAP, ALKBH5 and FTO) were depleted (Figure 1B).

**Figure 1.**
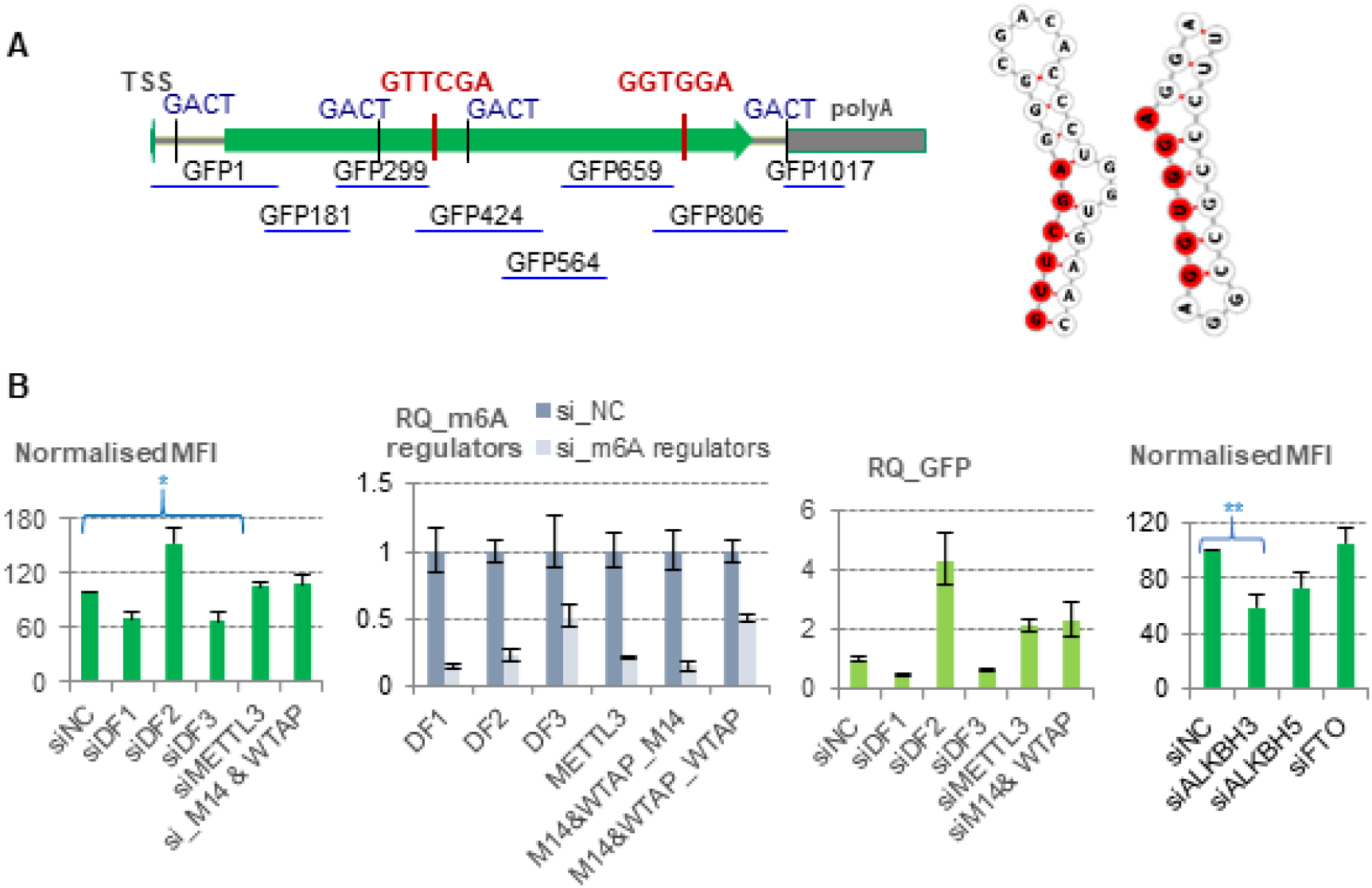
Impact of m6A and m1A regulators on GFP expression. **A.** Potential m6A (GACT) and m1A sites on GFP mRNA and the local RNA secondary structures of m1A sites using the RNA fold web server at http://rna.tbi.univie.ac.at/cgi-bin/RNAWebSuite/RNAfold.cgi (red and red highlighted letters are putative m1A sites). The overlapping regions amplified in qRTPCR (light blue lines) are shown. Transcriptional start site (TSS). **B.** Expression of GFP in CHO-K1 silenced for m6A writers and readers (left), m6A and m1A erasers (right) are shown as normalized Mean Fluorescence Intensity (MFI). Values are the means ± standard errors (SD), n = 3, p <=0.05 (*), p <= 0.01 (**), Student’s t-test (tails; 2 and type; 2). Relative expression (RQ) of m6A writers and readers in CHO-K1 silenced for m6A writers and readers (second left) and GFP (third left) determined by qRT-PCR. siRNA for negative control (siNC), m6A writer METTL3 (siMETTL3), m6A writers METTL14 and WTAP (siM14&WTAP), DF readers (siDF1, siDF2 and siDF3), m6A erasers (siALKBH5 and siFTO), m1A eraser (siALKBH3). RQ of METTL14, WTAP in CHO-K1 transfected with siRNA for both METTL14 and WTAP (M14&WTAP_M14, M14&WTAP_WTAP).

### Interplay between m6A reader DF2 and m1A eraser in regulation of GFP expression

It has been suggested that members of the DF sub-family and YTHDC1 can bind directly to m1A in RNA. In particular, the conserved Trp^432^ in the YTH domain of DF2 (Figure S1), which is necessary for its binding to m6A, is required for its recognition of m1A (31). We therefore investigated if there was cross-talk between DF2 and ALKBH3 in regulating GFP expression. We characterized the expression of GFP in CHO-K1 cells over-expressing DF2 and knocked-down for either ALKBH3, ALKBH5 or FTO. Over-expression of DF2 (CMV-DF2) reduced the expression of GFP in cells co-transfected with the control siNC, as expected. When co-transfected with siALKBH3 however, the expression of GFP was further reduced (Figure 2A), while no further changes in GFP were observed in cells transfected with both CMV-DF2 and siALKBH5 or siFTO, suggesting that DF2, in addition to acting as a m6A reader, most likely also binds the m1A target site of ALKBH3 to increase the negative impact of siALKBH3 by inducing further degradation of GFP. Over-expression of the human ALKBH3 which shares more than 93% amino acid identity with its CHO counterpart did not enhance the level of GFP produced in wild-type CHO-K1as expected. Therefore, in cells co-transfected with both siDF2 and the human ALKBH3 over-expression vector, the combined effect on GFP was unchanged compared with that resulting from the knockdown of DF2 (Figure 2B).

**Figure 2.**
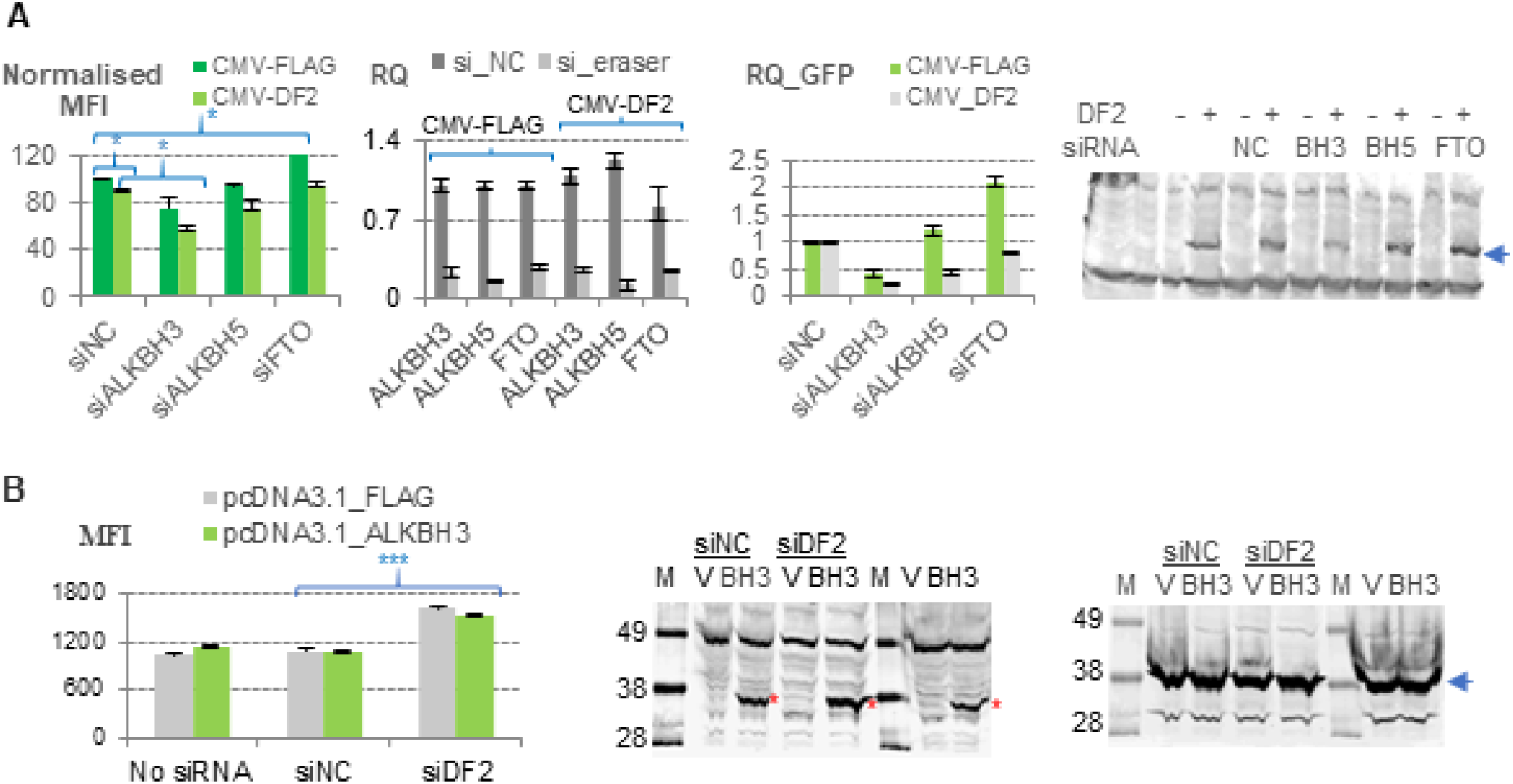
Impact of knockdown of erasers and over-expression of DF2 on GFP expression. **A.** Expression of GFP protein in CHO-K1 cells co-transfected with siRNA and CMV-DF2 is shown as MFI (left). Values are the means ± standard errors, n=2, p <=0.05 (*), Student’s t-test (tails; 2 and type; 2). Relative abundance of m6A and m1A erasers (second left) and GFP mRNA (third left), in CHO-K1 transfected with siRNA and CMV-DF2 determined by qRTPCR. Over-expression of DF2 (CMV-DF2) determined by Western blot analysis using an anti-FLAG antibody (right); blue arrow on the right is the expected molecular weight (65 kilo Daltons) of DF2 open reading frame and 2 FLAG epitopes; Negative control (NC), ALKBH3 (BH3), ALKBH5 (BH5); FTO (FTO). **B.** Effects of overexpression of ALKBH3 and silencing of DF2 on the expression of GFP. Expression of GFP in CHO-K1 co-transfected with human-ALKBH3 vector and siRNA measured by MFI (left). Values are the means ± standard errors, n=2, p <=0.001 (***), Student’s t-test (tail, 2; type; 2). Western blot analysis using an anti-FLAG antibody (middle) or an anti-GAPDH (right). Protein marker (M) with molecular weights are shown; pcDNA3-FLAG (V); pcDNA3.1-ALKBH3 (BH3). Negative control (NC). The position of overexpressed human ALKBH3 protein and endogenous gapdh are marked by red stars and blue arrow, respectively.

### GFP mRNA is immunoprecipitated with an anti-m6A antibody

We then performed m6A immunoprecipitation with an anti-m6A antibody using RNA prepared from CHO-K1 cells expressing the GFP construct and treated with siNC or siDF2. The m6A_RNA control was successfully precipitated using the anti-m6A antibody as expected, unlike the unmodified RNA (Figure 3A). In CHO-K1 cells transfected with siRNA (NC or DF2) and GFP, GFP mRNA was enriched in the m6A_IP fraction compared with the Input (Figure 3C). Similarly, positive controls SON and METTL14 mRNA which are known to be methylated (7, 25) were pulled down by the anti-m6A antibody. The percentage of m6A_IP/Input for GFP, SON, and METTL14 were similar (3-7%) while no significant enrichment was observed in the control gapdh_IP/Input (%) (Figure 3B). The ribosomal protein L30 (RPL30) mRNA, on the other hand, was not enriched and therefore not subjected to m6A methylation as expected (4).

**Figure 3:**
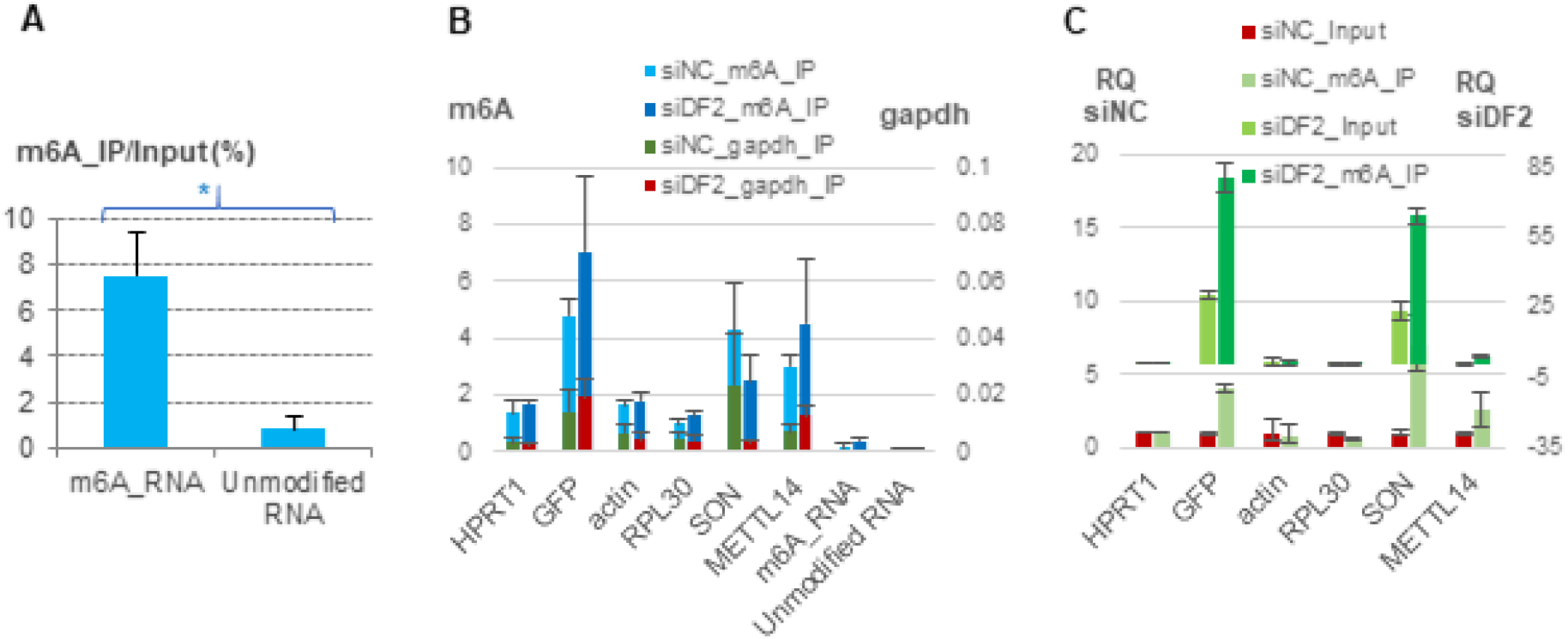
GFP mRNA is immunoprecipitated by an anti-m6A antibody (n=3) Percentage of m6A_IP/Input of control mRNAs (A), mRNA from CHO-K1 co-transfected with of siRNA and GFP immunoprecipitated using an anti-m6A antibody or anti-gapdh antibody (B). Values represent the mean ± standard error (SD), p <=0.05 (*), Student’s t-test (tails; 2 and type; 2). Relative expression of GFP in m6A_IP and Input fractions determined by qRTPCR (C). Immunoprecipitate (IP).

### GFP mRNA is immunoprecipitated with an anti-m1A antibody

As our genetic data indicated the presence of m1A on the GFP transcript, we performed immunoprecipitation with an anti-m1A antibody using mRNA from cells transfected with either siNC or siALKBH3 and the GFP expression construct. We found that the well-characterized m1A site at nucleotide 1322 in human 28S rRNA, which is known to be conserved between mouse and human, is also conserved in CHO-K1 and is located at nucleotide position 1146 (GenBank accession number NR_045212) (Figure 4A). As shown in Figure 4B, an RT-PCR of 28S rRNA amplified using primers spanning the m1A site (28S) was not as efficient (average Ct 21.4) as those amplifying a region upstream (28S_304) (average Ct 10.89) or downstream (28S_1332) (average Ct 11.1), due to its position at the Watson-Crick interface, which effectively stalls reverse transcription (RT) resulting in truncated RT products (32). The anti-m1A antibody did not capture the spiked m6A_RNA or unmodified RNA, as expected. However, the GFP transcript was effectively precipitated with the anti-m1A antibody, as demonstrated using a panel of primers against different regions of the mRNA, with a ratio of m1A_IP/Input between 0.5 to 1.2% (Figure 4C, left panel). The m1A_IP/Input of 28S, which is known to be highly m1A methylated (23, 24), ranged from 1.2 to 2.9%. In addition, we found that the ALKBH3 transcript which has two putative m1A sites GTTCAA at bases 379 and 550 (Figure 4D) was also immunoprecipitated. An RT-PCR amplified using primer pairs spanning the two putative m1A sites (BH3_320 and BH3_470) were significantly less efficient, compared to that of a region downstream (BH3), although the difference was not as striking as for the 28S rRNA. This was observed only in input mRNA from cells transfected with siNC control but not with siALKBH3 (Figure 4D). We reasoned that in cells with ALKBH3 deleted, the m1A sites could be more accessible to endogenous DF2 which accelerates degradation of m1A containing mRNA. Despite scanning the entire GFP transcript from the transcriptional start site to the polyA (Figure 1A) using qRT-PCR, we did not identify the RT-arrest region, the signature of the m1A site on RNA. It is possible that the m1A modification on GFP is low or the methylation is incomplete, so there would not be enough difference between the amplicon spanning the m1A site and the rest of the GFP transcript, as in the case of 28S rRNA 1322 m1A site (more than 70% methylation in human KEK293T cells) (24). GFP levels determined by RT-PCR was lower in the Input fraction of CHO-K1 silenced for ALKBH3, suggesting ALKBH3 affects GFP at the transcriptional level. We found that the level of 28S rRNA increased in the m1A_IP fraction of CHOK1 transfected with siALKBH3 (Figure 4D, right panel). This is consistent with its role as a m1A eraser (21, 22). However, the level of GFP and ALKBH3 in cells depleted of ALKBH3 in the m1A_IP fractions were reduced, in contrast with that of 28S rRNA (Figure 4C, right panel). It appears that the fate of m1A containing ALKBH3 and GFP mRNA was different from that of 28S rRNA in cells depleted for ALKBH3. The mechanism underlying the difference is yet to be determined. Nucleomethylin has been shown to be required for the N1-methyladenosine (m1A) modification in 28S rRNAs whose m1A site is GTTCGA in human and mouse cells (33), while tRNA methylase complex TRMT6/61A has been implicated in the m1A methylation of a small subset of mRNAs at m1A sites with t-RNA T-loop-like structures outside the 5’UTR (23, 24). It remains to be determined the writer responsible for installing m1A in GFP/ALKBH3 mRNA.

**Figure 4:**
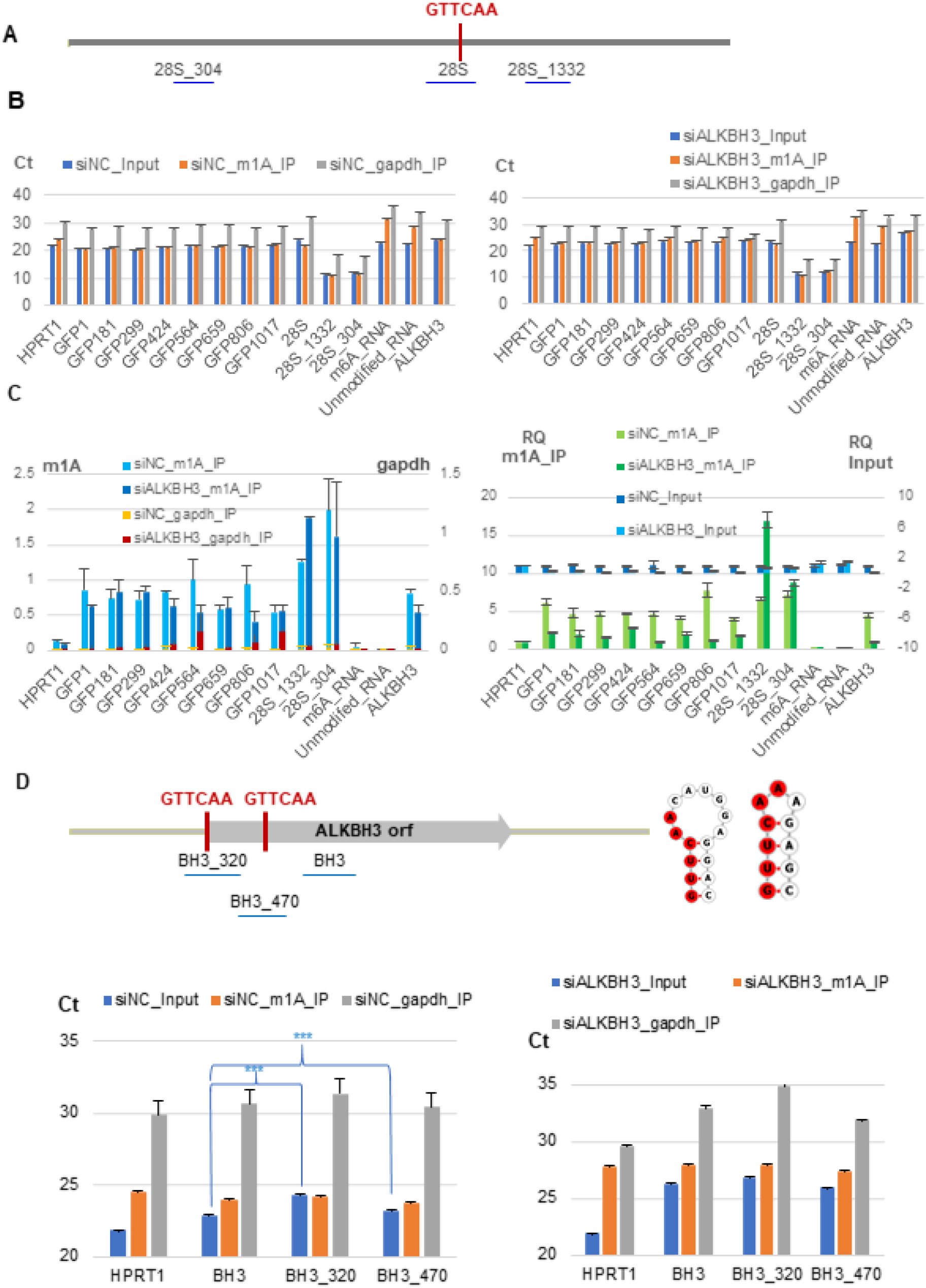
GFP mRNA is immunoprecipitated by an anti-m1A-antibody (n=2) **A.** Conserved m1A site on CHO 28S rRNA (GenBank accession number NR_045212) (red letters) and regions used in qRTPCR (light blue lines) are shown. **B.** Relative gene expression represented as Ct values after immunoprecipitation using an anti-m1A antibody with mRNA from CHO-K1 co-transfected with GFP vector and siNC (left) or siALKBH3 (right). Immunoprecipitate (IP). **C.** Percentage of m1A_IP/Input of mRNA from CHO-K1 co-transfected with siRNA and GFP immunoprecipitated using an anti-m1A antibody (primary axis) or anti-gapdh antibody (secondary axis) (left panel). Values are the means ± standard errors. Relative expression of GFP in m1A_IP and Input fractions determined by qRT-PCR (right panel). Immunoprecipitate (IP). **D.** Putative m1A sites on ALKBH3 mRNA of CHO-K1 (GenBank accession number XM_007648356) and their local secondary structures (upper panel). The first putative m1A site is at - 2 relative to ATG, GGTCAA in human ALKBH3 (NM_139178); the second site is conserved between the two species. Regions used in qRTPCR (light blue lines) are shown. Relative gene expression of ALKBH3 amplified with different primer pairs represented as Ct values after immunoprecipitation using an anti-m1A antibody with mRNA from CHO-K1 co-transfected with GFP vector and siNC or siALKBH3 (left and right, bottom panel). Immunoprecipitate (IP). p<= 0.001 (***). Student’s t-test (tails; 2 and type; 2).

Research in the last few years has highlighted the variable nature of the prevalent m6A modification and its impact on diverse biological processes (34–36). A few recent studies have implicated that the m1A mark is also a dynamic modification responsive to changing physiological and environmental conditions (21, 22) such as mammalian cells are exposed to in a bioreactor. These modifications eventually affect mRNA fate and translation efficiency-potentially impacting the production of biopharmaceutical proteins from cells. This study provides the first evidence of the significance of the m6A readers DF1-3 on the expression of a model transgene, GFP. In particular, silencing of DF2, the main m6A reader that regulates mRNA stability has a pronounced impact on protein expression in CHO cells. A similar impact of DF2 knockdown was also observed on expression of human Erythropoietin, a drug used for the treatment of some forms of anemia. This study is also the first to show that the expression of an mRNA can also be subjected to co-ordinated modulation of both m6A and m1A regulators. We also present the first genetic data implying that the adenosine nucleotides targeted for m1A methylation are read by a m6A reader protein. In addition, we identified that the m1A eraser ALKBH3 itself is also a target for m1A modification in CHO cells. To our knowledge, although methylation of the ALKBH3 mRNA was found to be located at different sites in m1A transcriptomic analysis of human liver carcinoma cells subjected to heat stress, it was not identified in HEK293 cells, Hela cells or various mouse cell types (21–24). The two potential m1A sites in human and CHO ALKBH3 mRNA are conserved in sequence and location. M1A sites in the 5’UTR have been found to be positively correlated with protein production (16, 17) while those in the coding region were not (23, 24). We demonstrated that the ALKBH3 mRNA is precipitated by an anti-m1A antibody though is yet to be determined which sites are targets for m1A methylation and their exact roles.

At present, our knowledge of the nature and exact role of the m1A methylome in mammalian cells is limited, compared to that of the m6A, partly due to the technical challenge of single m1A nucleotide resolution mapping (37). Our findings enrich and clarify some ambiguities in our current knowledge of m6A and m1A regulation in mammalian cells in general. In particular, these insights open the prospect of new epi-transcriptomic-based approaches to mammalian cell line engineering for recombinant protein production. An epi-transcriptomic-based approach could be developed based on insight gained from this study to enhance transgene expression in mammalian expression systems, e.g. development of a cell line with different level of DF2 expression as a host for production of biopharmaceutical proteins or incorporation of m1A sites in the 5’UTR of a recombinant gene sequence. Given the fact that the m6A reader DF2 and eraser ALKBH3 have opposite and significant impact on the level of the transgene GFP mRNA, identification of the endogenous mRNA targets of these two RNA methylation regulators is the logical next step to understanding the cross-talk between m6A and m1A networks at cellular level in general or during bioproduction processes in CHO in particular. Finally, this study demonstrates that identifying the impact of and potentially manipulating the methylation events on product-encoding sequences could form the basis for more efficient therapeutic protein production strategies.

## FUNDING

This work was supported by the Scientific Foundation Ireland [13/IA/1963 to N.B.]. Funding for open access charge: Scientific Foundation Ireland.

## CONFLICT OF INTEREST

The authors declare no competing interest

