## Supplemental information for "Cross-talk between m6A and m1A regulators, YTHDF2 and ALKBH3 fine-tunes mRNA expression"

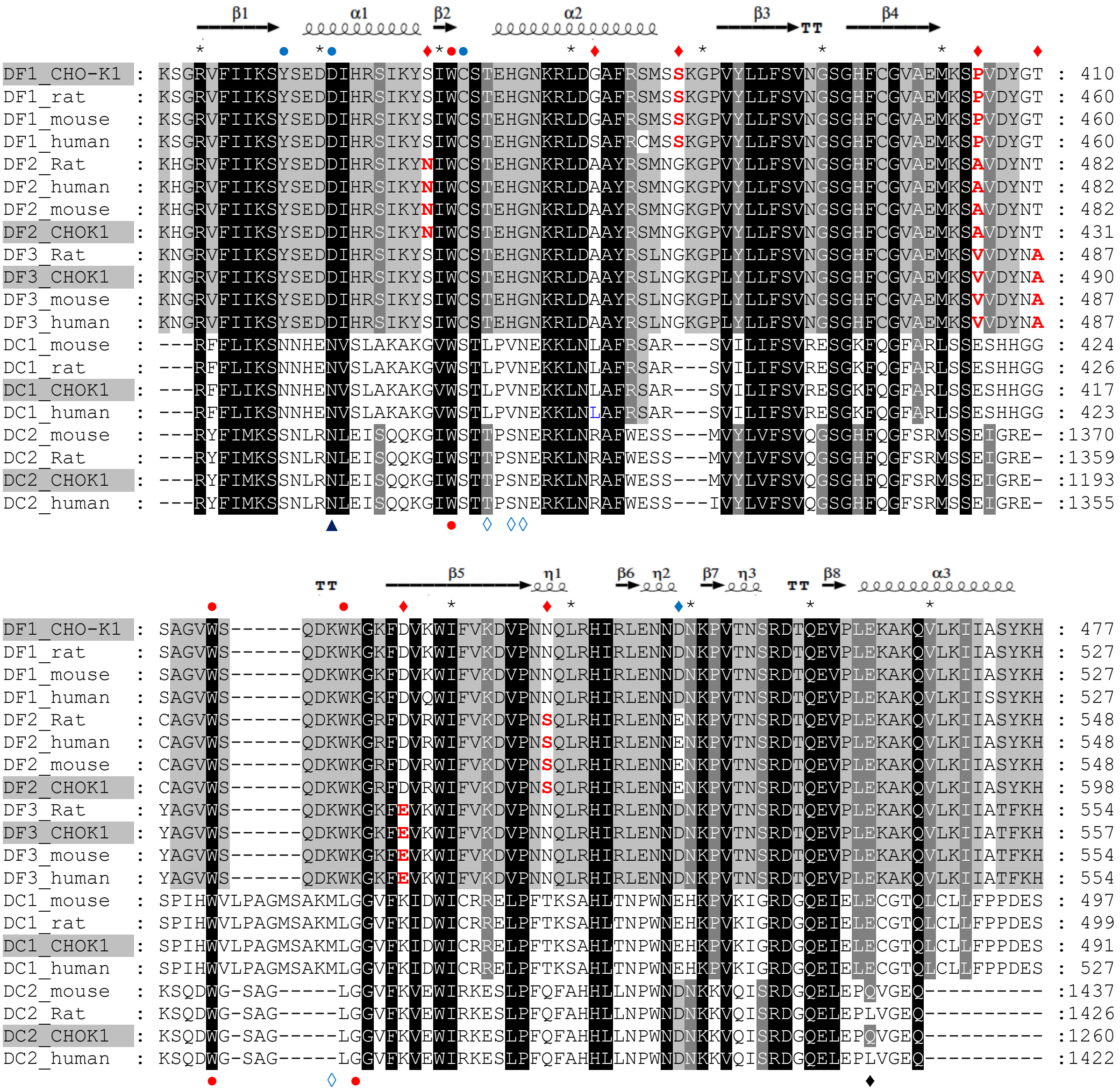


**Figure S1**. Multiple alignment of YTH domains

YTH domain amino acid sequences of YTH containing proteins from mouse, rat, human and CHO cells were aligned using CLUSTALX software with default settings. Identical or similar residues shared by different number of sequences were displayed using GenDocs program with shading from white to black. Secondary elements in the YTH domain shown above the alignment were from the on-line service at http://espript.ibcp.fr/ESPript/ESPript/ using the protein structure of the YTH domain of human DF1 (PDB: 4RCJ). Amino acid positions are shown at the end of each line on the right side. TT, strict β-turns; arrows, β-strands; coils, α-helices; η, $3_{10}$-helix. M6A pockets of DF and DC1 are shown above and below the alignment, respectively. Red, full circles, aromatic residues critical for m6A recognition (WWW in DF proteins and WWL in DC proteins); blue shuttle shape, residues that are DC1 unique for the preference of G-1 (the position preceding the m6A nucleotide); dark blue triangle, residues has been experimentally shown to involve in the stronger m6A binding activity of the YTH domain of DC1; dark blue shuttle shape under the alignment, residues unique for CHO-K1 and mouse in the YTH domain of DC2; red filled shuttle above the alignment, residues unique for either DF1, DF2 or DF3; sky-blue filled shuttle, residues unique for DF2 and DC1.

**Figure S2**. Impact of knockdown DF2 and FTO on EPO expression.

Western blot analysis of EPO expression in CHO-K1 cells co-transfected with siRNA (top) and quantification (bottom). SiRNA for negative control (NC). Molecular weight (M), Standard EPO in ng (1, 52). Samples taken for analysis: 30uL supernatant at Day 2 post transfection. Blue arrow on the left is the expected molecular weight (38kD) of glycosylated EPO.

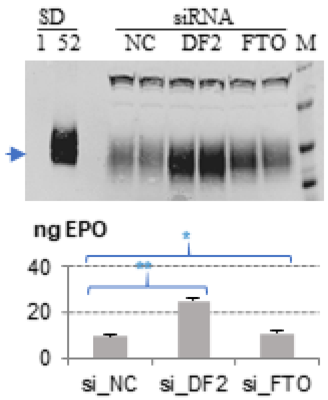


**Table S1**

A. **Primers sequences** (5'-3') used in the construction of overexpression vector CMV-DF2

| Primers | Sequence **(5'-3')** |
|---|---|
| DF2for | ATGTCGGCCAGCAGCCTCTTGGA |
| DF2rev | TTATTTCCCACGGCCTTGACGCTCCTTTTTAACA |
| FLAG2YN | ACAAGGATCATGATATTGATTACAAGGACGACGATGACAAGATGTC GGCCAGCA |
| 3FLAGBamfor | GGGGATCCATGGACTATAAGGACCACGACGGAGACTACAAGGATCA TGAT |
| Y2endXho | GATCTCGAGTTATTTCCCACGGCCTTGACGCT |

B. **SiRNA sequences (5'-3')**

| Gene | Si_RNASequence (5'-3') |
|---|---|
| DF1 | CAAATGTAAACATGCCAGTTTCA |
| DF2 | ACCTAAACTGAAGACCAAGAATGGC |
| DF3 | CTACCATTGGTGCAAAGCCAACTGC |
| METTL3 | AAATTGATGCTTGCATGGATTCTGAG |
| METTL14 | ACAAAGATTCCAGTACCTTTCTTAA |
| WTAP | GCAGTCAGGATGAACTGAATGAC |
| ALKBH3 | CCCCATCATTGCGTCACTTAGTTTT |
| FTO | CACTTGGCATGTTGGTTTCAAGATA |

C. **Primers for qPCR**

| **Genes** | **Forward primers (5' -3')** | **Reverse primers (5' -3')** |
|---|---|---|
| DF1 | CAGTTAAGACGGTGGGTTCAG | CATCTTGGGCTGTGGTTTTG |
| DF2 | CAGTTTGCCTCCAGCTACTATT | GCAAGGCCATTCTTGGTCTTC |

| | | |
|---|---|---|
| DF3 | AGCAGTGGTATGACTAGCATTG | CAATTCCCACATTGCCCTTG |
| METTL3 | TTGCATGGATTCTGAGGGTC | CCAGGTAGCGGATATCACAAC |
| METTL14 | CGGAAGTGCCTGGATGAAG | AAGCTCTGCGTTCCCTTAAG |
| WTAP | AGCAGAGTTGGCTTTACAGAAG | CTGTCGTGTCTCCTTCAATTG |
| ALKBH3-human | TGTATCCTGGCTTTGTTGAC | TCTCCATACCATGCTGTAAGTC |
| **ALKBH3-CHO** | TGTCTATATCCTGGCTTTGTGG | GCTCTCCATACCATGCTGTAAG |
| BH3_320 | CATGGCTTGAATTGGGAATGA | TTCCTAGCAGCAGTACCTGGCT |
| BH3_470 | TAGGAACCATCTTTCCCAGACA | TGTCAATCACTCTGGGTTCTG |
| HPRT1 | TTTACCTCACCGCTTTCTCG | TCATCACTAATCACGACGCTG |
| SON | CTGTAACAGTAGGAGTGGATC | GGAGTCCATAGTGCTAGAGGC |
| RPL30 | GTCCATCAACTCGAGGCTC | TTTCAGATTTCCTCAGGGCC |
| FTO | TGCACCATCAATTACACAGAGG | CACAAAGGCACAGCATCTTC |
| 28S | GTTCTCTCCGGTCACGC | TGACTCGCGCACGCGTTAGAC |
| 28S_304 | CAAAGCGGGTGGTAAACTCC | TTCACGCCCTCTTGAACTCT |
| 28S_1322 | GCGTTAGGACCCGAAAGATG | GTCTTTCGCCCCTATACCCA |
| M6A_RNA | CGACATTCCTGAGATTCCTGG | TTGAGCAGGTCAGAACACTG |
| Unmodified RNA | CTGAAGGAGCCTGTGATCTG | GTGGCACACGTTACATTTCTG |
| EPO | TGTGGATAAAGCCGTCAGTG | GGAAGAGTTTGCGGAAAGTG |
| GFP1 | GAGAGAACCCACTGCTTACT | GAACTTGTGGCCGTTTACGT |
| GFP181 | GACGTAAACGGCCACAAGTT | GTAGGTCAGGGTGGTCACGA |
| GFP299 | TCGTGACCACCCTGACCTAC | GTCTTGTAGTTGCCGTCGTC |
| GFP424 | GACGACGGCAACTACAAGAC | GGCGGATCTTGAAGTTCACC |
| **GFP564** | ACAACGTCTATATCATGGCCG | GTGCTCAGGTAGTGGTTGTC |
| GFP659 | ACCACTACCAGCAGAACACC | CTTGTACAGCTCGTCCATGC |
| GFP806 | ATCACTCTCGGCATGGACGA | TGGCAACTAGAAGGCACAGT |
| GFP1017 | CGACTGTGCCTTCTAGTTGC | AGGAAAGGACAGTGGGAGTG |

**Table S1**. Primers used in the construction of the vector CMV-DF2 (A), siRNA sequences (B) and in qPCR (C).

Regular used primers for qRTPCR of GFP and ALKBH3 (BH3) are bold.
